# Evaluation of methods for AlphaFold-based integrative modeling

**DOI:** 10.64898/2026.09.13.751323

**Authors:** Kartik Majila, Shruthi Viswanath

**Affiliations:** National Centre for Biological Sciences, Tata Institute of Fundamental Research, Bangalore, Karnataka, India 560065

**Keywords:** AlphaFold, Integrative modeling, Macromolecular assemblies, Structure prediction, Chemical crosslinking

## Abstract

Recent methods enable incorporation of experimental data, such as chemical crosslinks, X- ray and cryo-EM density maps, into AlphaFold to guide the predictions towards structures consistent with the input data. However, it is unclear how these methods balance the input data with AlphaFold’s learned structural priors. To probe this aspect, we curated a benchmark of homomeric and heteromeric protein complexes non-redundant with the AlphaFold training set. We evaluated three methods, AlphaLink2, Boltz2, and GRASP, on the benchmark, using chemical crosslinks as input data. All methods predict structures that satisfy the crosslinks well, in the easy case, where 90% of the input crosslinks are consistent with a single structure. Yet this apparent success belies important limitations. First, for most complexes, the methods produce nearly identical predictions with and without the input crosslinks. Second, predictions remain largely unchanged from the easy case even as the fraction of false-positive crosslinks increases, even though satisfying these crosslinks would require sampling conformations that differ substantially from those predicted in the easy case. Finally, when challenged with input crosslinks from multiple states, the methods remain biased toward a single state. Together, these findings suggest that current AlphaFold-based approaches can be insufficiently sensitive to the input data, with the predictions likely driven primarily by the learned structural priors. Our study underscores the need for AF-based IM approaches that can weigh experimental evidence better and can capture structural heterogeneity. The benchmark and all code to obtain the predictions and perform the analysis are available at https://github.com/isblab/af_im.

## Introduction

Macromolecular complexes play important roles in various cellular processes, including transcription, translation, molecular trafficking, and signal transduction, yet characterizing their three-dimensional structures remains challenging ^1,2^. Often, experimental structure determination techniques like X-ray crystallography, nuclear magnetic resonance (NMR), and cryo-electron microscopy (cryo-EM) remain limited due to challenges in sample preparation and the size and heterogeneity of the complexes ^1,3^. Integrative modeling (IM) is an approach that overcomes the limitations of individual techniques by integrating data from various experimental techniques along with physical principles, statistical inferences, and prior models ^1,4–7^. Given sparse, noisy, and ambiguous input data from heterogeneous sources, no single structure may satisfy all the data. As a result, the aim of integrative modeling is to determine the set of all structures consistent with the input data ^1,3,7^.

Advancements in deep learning methods like AlphaFold (AF) have transformed structural biology, enabling accurate sequence-based structure prediction ^8–10^. Although methods such as AlphaFold-multimer and AlphaFold3 enable modeling complexes, they are often inaccurate on challenging targets, including large complexes, disordered proteins, and multi-state proteins ^11–14^. Recent methodological advances have led to the development of AlphaFold-based integrative modeling (AF- based IM) methods that allow data-guided structure prediction ^13^. Examples include AlphaLink/AlphaLink2, GRASP, Distance-AF, ColabDock, ROCKET, Guided-AF3, ConforNets, and Boltz2 ^13,15–22^. In contrast to methods that use AlphaFold predictions as input for downstream modeling, AF-based IM methods allow incorporating experimental data to influence the AlphaFold predictions ^13^. The experimental data is incorporated in various ways: as a bias for the MSA and pair representations, as restraints in the loss function for fine-tuning, or as a bias in the diffusion sampling process ^13^. Specifically, AlphaLink/AlphaLink2 incorporates crosslinking data into the pair representation used as input to AlphaFold’s Evoformer ^19,20^. Similarly, GRASP incorporates residue-level interaction data into the MSA representation input to the Evoformer and the single representation input to the structure module ^21^. In contrast, Boltz2 incorporates distance restraints from experimental data to bias the diffusion sampling at inference time. Thus, by combining AlphaFold’s learned structural priors with biophysical information, these methods potentially enable data-guided conformational search ^18^.

Despite the rapid progress of these AF-based IM methods, their applicability and limitations have not been systematically evaluated. It remains unknown how these methods weigh the learned structural priors with experimental restraints, particularly when dealing with noisy data and data derived from multiple states. In this study, we benchmark the performance of state-of-the-art AF-based IM methods – AlphaLink2, Boltz2, and GRASP – on a dataset of 37 multi-protein complexes using chemical crosslinks as input. We assess the ability of these methods to predict structures consistent with the input crosslinks. We evaluate the set of predicted structures from these methods based on their accuracy, stereochemical plausibility, and diversity. We also test the robustness of these methods to noise in the crosslinking data. Finally, we probe their ability to predict distinct states based on crosslinks from multiple states. Our findings suggest that current AlphaFold-based approaches can be insufficiently sensitive to the input data, with the predictions likely driven primarily by the learned structural priors. Overall, our study highlights several limitations of the AF-based IM methods and points to directions for future improvements.

## Results

To assess the performance of the AF-based integrative modeling methods, we created a benchmark of 37 complexes non-redundant with the AlphaFold training set (Fig. 1, Suppl Table 1, see Methods). Input crosslinks were simulated from the known experimental structure (native structure) of each complex using JWalk ^32^. Ninety percent of these crosslinks were consistent with the native structure (SASD≤20 Å; true positives), whereas the remaining 10% were adversarial crosslinks inconsistent with the native structure (SASD>40 Å; false positives), reflecting the false positive rate observed in crosslinking experiments (Fig. 1). Three AF-based IM methods were evaluated: AlphaLink, Boltz, and GRASP, with twenty-five structures predicted by each method per complex.

**Figure 1:**
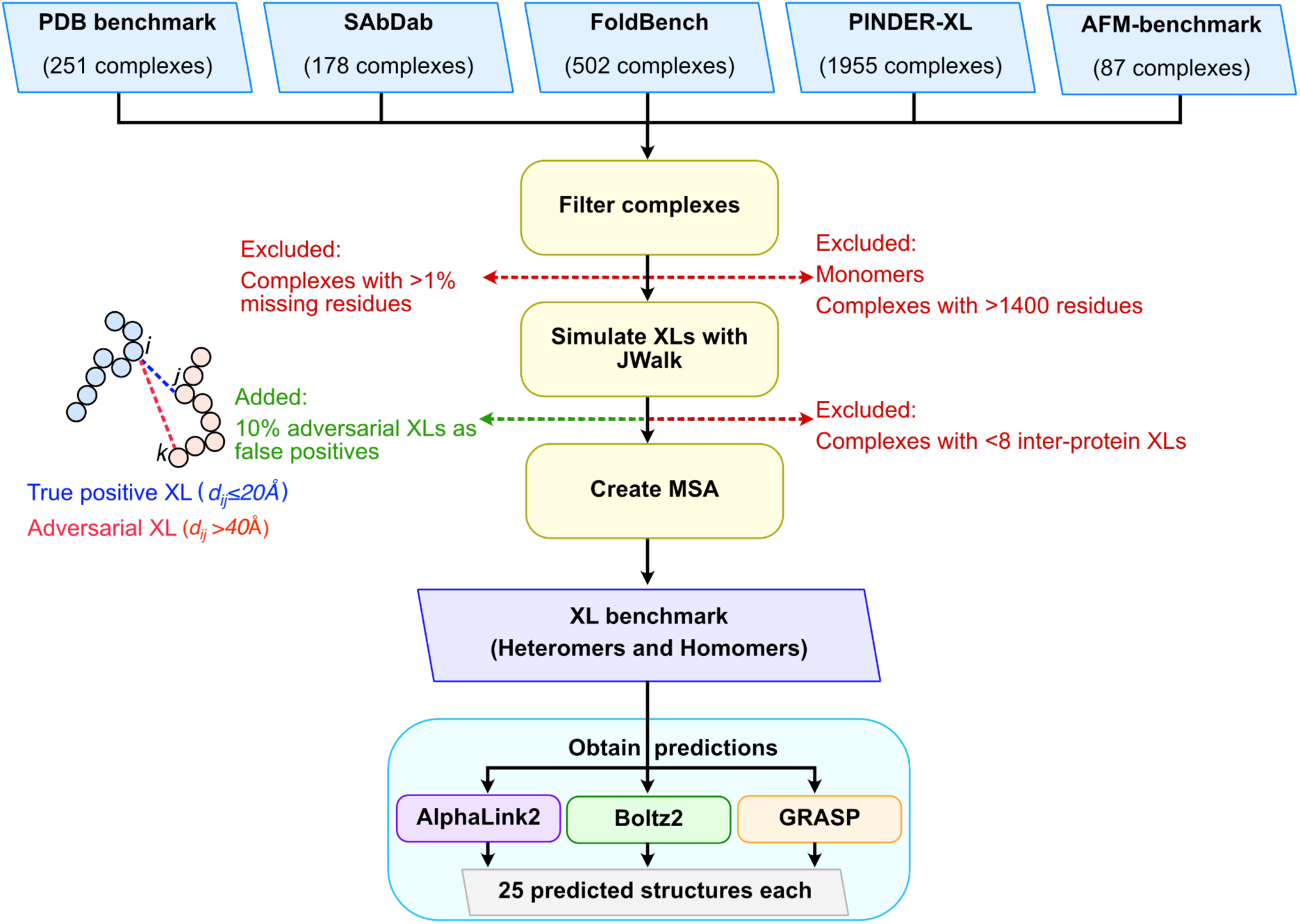
Dataset creation pipeline. Our dataset was compiled by gathering protein complexes from existing benchmarks, including the PDB benchmark, SAbDab, FoldBench, PINDER-XL, and AFM-benchmark. Monomeric proteins, complexes with missing residues, and those greater than 1400 residues in length were excluded. Inter- protein crosslinks (XLs) for each complex were computed using JWalk, and complexes with fewer than eight crosslinks were excluded. Residue pairs with a solvent accessible surface distance (SASD) less than 20 Å were considered as true positive (TP) crosslinks. Additionally, 10% adversarial false positive (FP) crosslinks with a SASD greater than 40 Å were used. MSAs for each complex were computed using the OpenFold MSA creation pipeline. Three AF-based IM methods were selected – AlphaLink2, Boltz2, and GRASP for evaluation.

### Do the predicted structures satisfy the data?

First, we assessed whether the AF-based IM methods could predict structures that satisfied the input crosslinks. The crosslink satisfaction for a predicted structure is the fraction of crosslinks satisfied by the structure. We measured the maximum crosslink satisfaction achieved across all predicted structures from each method for each complex. For most complexes, all the methods predicted at least one structure with high crosslink satisfaction, *i.e.,* more than 75% crosslinks satisfied (Fig. 2). GRASP predicted structures with high crosslink satisfaction for all the complexes, whereas AlphaLink2 and Boltz2 predicted structures with high crosslink satisfaction for 35 and 34 complexes, respectively (Fig. 2, Supp Table 1).

**Figure 2:**
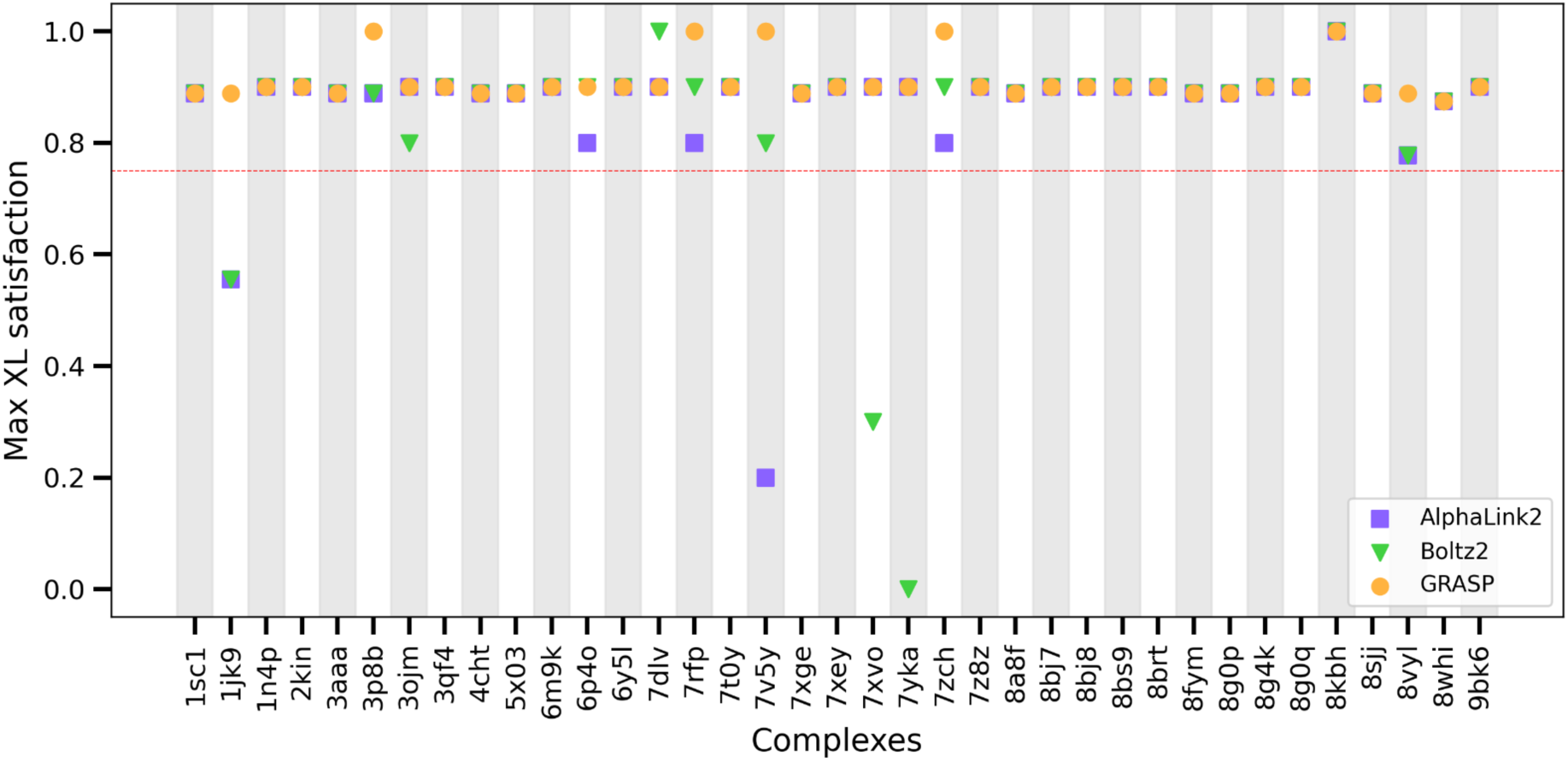
**Assessing crosslink satisfaction**. For each complex, the maximum crosslink (XL) satisfaction achieved by a structure in the set of predicted structures is shown for AlphaLink2 (purple, squares), Boltz2 (green, triangles), and GRASP (orange, circles).

### Do guided predictions improve data satisfaction?

Having established that the AF-based IM methods could predict structures consistent with the input crosslinks, we next examined whether using crosslinks to guide the prediction improves crosslink satisfaction relative to unguided predictions, in which no crosslinks are provided. For this, we compared Boltz2 and GRASP, since they allowed performing unguided predictions, whereas AlphaLink2 does not. Comparing the distribution of per-model crosslink satisfaction between guided and unguided predictions, we observed that crosslink guidance improves crosslink satisfaction for GRASP (Supp Fig. 1, 2A). However, for Boltz2, the per-model distributions were nearly identical for both guided and unguided predictions, potentially suggesting a strong training bias (Supp Fig. 1, 2B).

### Are the predicted structures close to the native state?

High crosslink satisfaction alone does not imply structural accuracy, *i.e.,* similarity of the predicted structures to the native structure. We quantified structural accuracy using the maximum DockQ score between a predicted structure and the corresponding native structure. DockQ measures the interface similarity between two structures ^23,24^. AlphaLink2, GRASP, and Boltz2 predicted at least one near-native structure, *i.e.*, a structure with an acceptable DockQ (DockQ > 0.23), for 36, 36, and 34 complexes, respectively (Fig. 3A, Supp Table 1) ^24^.

**Figure 3:**
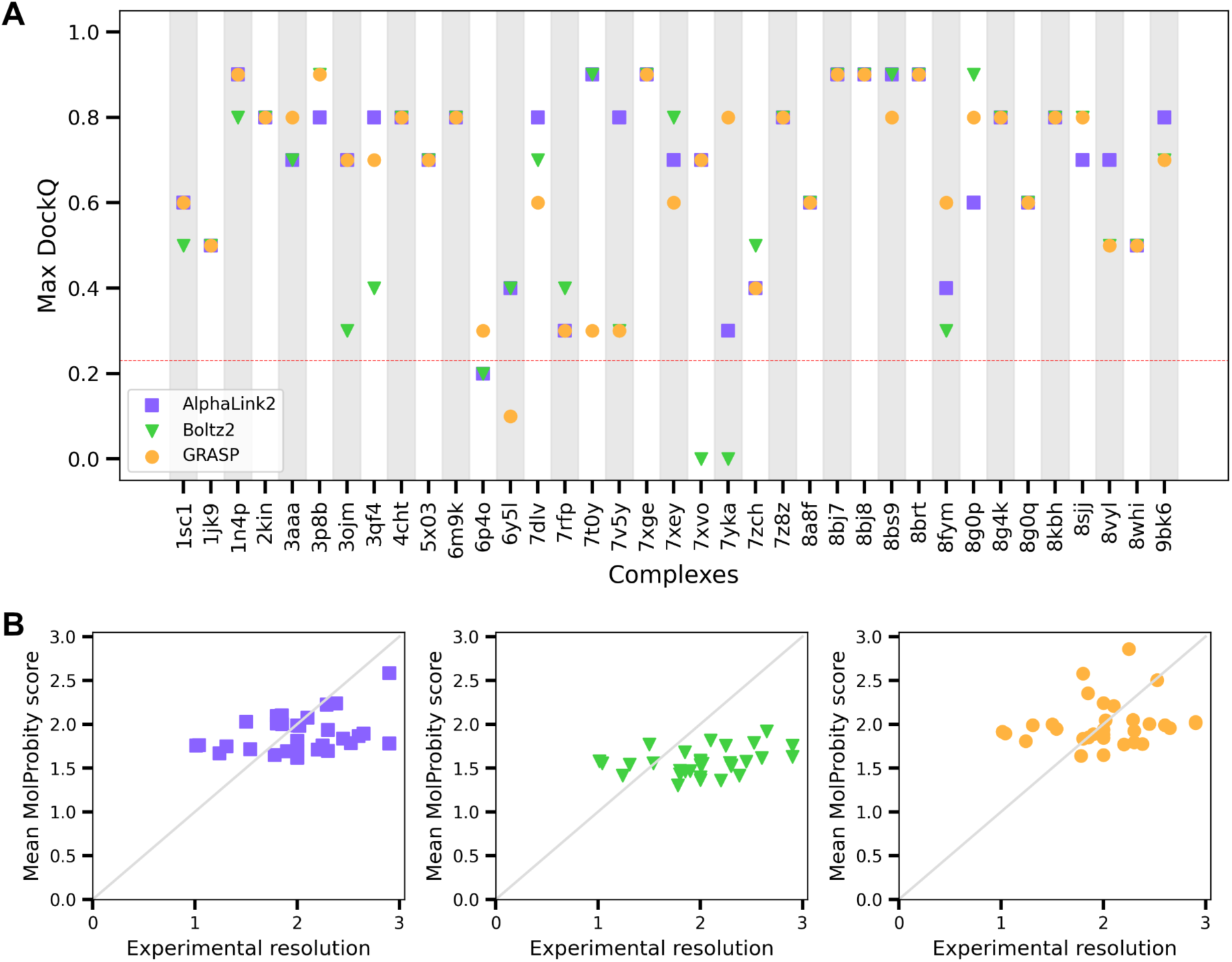
**Assessing structural accuracy and stereochemical plausibility**. For predictions obtained from AlphaLink2 (purple, squares), Boltz2 (green, triangles), and GRASP (orange, circles), (A) the maximum DockQ score achieved across all predicted structures for each complex is shown as a measure of structural accuracy, and (B) the mean MolProbity score across all predicted structures is shown as a function of the experimental resolution of the native structure.

### Are the predicted structures stereochemically plausible?

We further assessed the stereochemical plausibility of the predicted structures using MolProbity based on the MolProbity score and the clashscore ^25^. The MolProbity score reflects the crystallographic resolution of the structure. A predicted structure is better than the average structure at that resolution if the MolProbity score is lower than the crystallographic resolution. The clashscore indicates the number of inter-atomic clashes in the predicted structure, with a clashscore less than 5 considered ideal ^25^.

The mean MolProbity scores for the predicted structures by AlphaLink2, Boltz2, and GRASP were lower than the resolution of the experimental structure for 27, 31, and 24 complexes, respectively (Fig. 3B, Supp Table 1). This indicates that for most complexes, the predicted structures are stereochemically plausible. Of the three methods, GRASP predicted structures with the lowest stereochemical plausibility. We then compared the mean clashscore of the predicted structures from each method against the clashscore of the corresponding native structure (Supp Fig. 3). For all three methods, the mean clashscore across all complexes is similar to or higher than the clashscore for the native structure, indicating that the predicted structures have similar to or more inter-atomic clashes than the native structures. Clashscores from GRASP- predicted structures were higher than those from others for several complexes in the benchmark (Fig. 3).

### Is the set of predicted structures diverse?

Next, we assessed the methods based on their ability to sample a diverse set of structures, given input crosslinks. Given a sparse set of crosslinks with an upper bound of 20 Å, more than one structure may plausibly satisfy the data. To assess structural diversity, we first filtered the predicted structures based on interface similarity to obtain a set of unique structures (See Methods). We then quantified structural diversity by the number of unique structures predicted by each method for each complex. For most complexes, all three methods failed to predict a diverse set of structures (Fig. 4). GRASP predicted more than one structure for 11 complexes, whereas AlphaLink2 and Boltz2 did so for only 4 and 5 complexes, respectively, indicating that GRASP can sample a more diverse set of structures (Supp Table 1).

**Figure 4:**
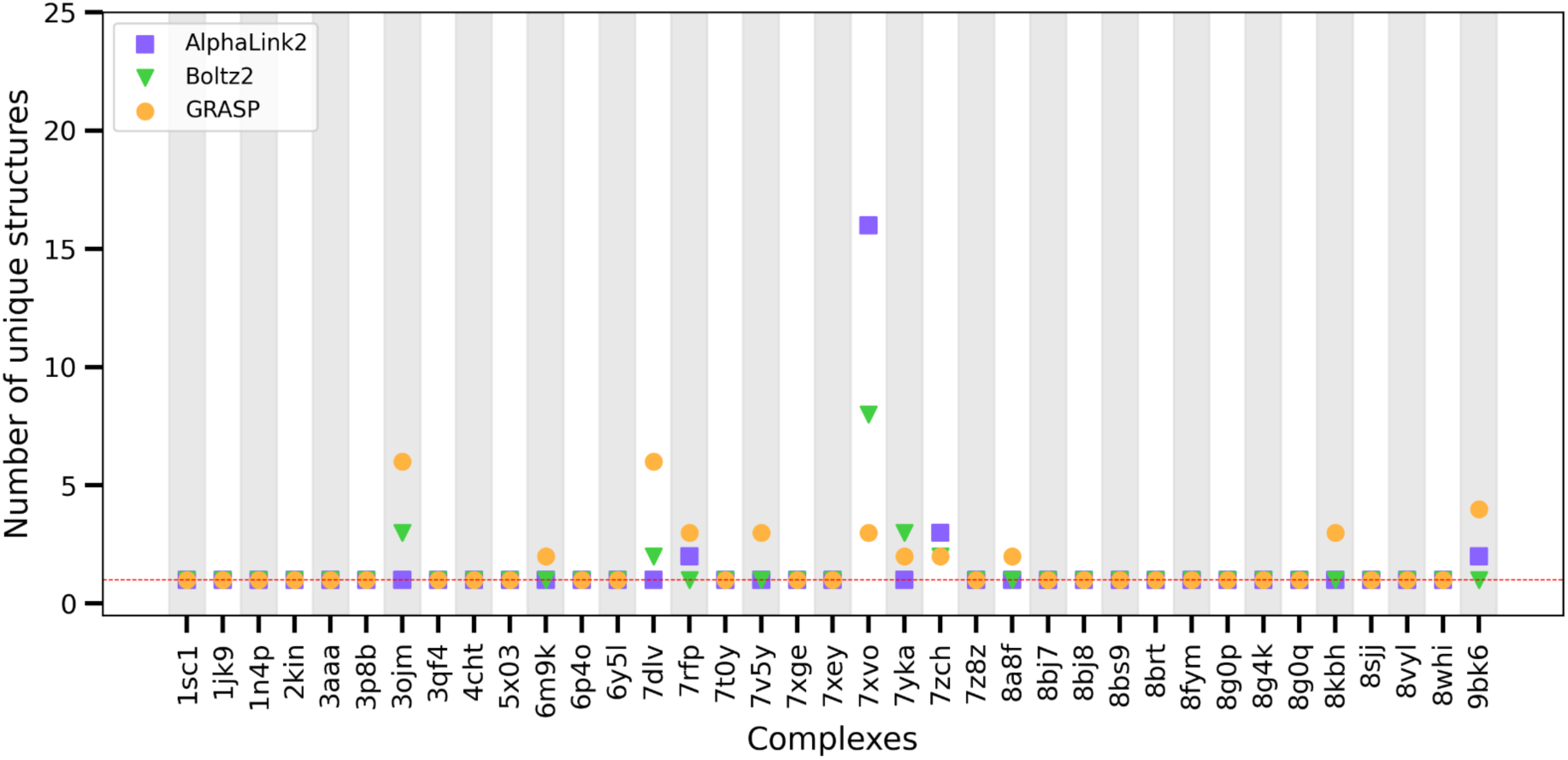
**Assessing structural diversity**. For each complex, the number of unique predicted structures by each method based on the DockQ score is shown for AlphaLink2 (purple, squares), Boltz2 (green, triangles), and GRASP (orange, circles).

We further probed whether the lack of diversity results from the model being highly confident in its predictions. For this, we measured mean model confidence across the set of predicted structures by each method across all complexes. The model confidence appears to be uncorrelated with the diversity of predicted structures (Supp Fig. 4). The Boltz2 (GRASP) predicted structures always have a high (low) confidence.

However, since most crosslinks used herein are derived from a single native structure per complex, there may not exist a diverse set of configurations consistent with the crosslinks for most complexes, despite the low resolution of crosslinking data. So, we cannot completely rule out the possibility that these methods may predict alternate configurations. Therefore, we proceeded to evaluate this aspect more directly, through two other assessments described in the next sections.

### Effect of false-positive crosslinks

Next, we evaluated the performance of the AF-based IM methods in response to a higher fraction (33%) of false-positive (FP) crosslinks. As noted previously, we used adversarial FP crosslinks for which the inter-residue distance exceeds 40 Å. Satisfying these FP crosslinks would require exploring configurations beyond the near-native structures; therefore, we expect the integrative modeling methods to predict a more diverse set of structures on this dataset.

For most complexes, all three methods satisfied the true-positive (TP) cross-links but failed to satisfy the false-positive (FP) cross-links, as measured by the maximum TP and FP cross-link satisfaction achieved by any structure (Fig. 5A,5B,5C). GRASP was slightly better at satisfying FP crosslinks, predicting structures that satisfy at least one FP crosslink for 14 complexes, whereas AlphaLink2 and Boltz2 satisfied FP crosslinks for only 6 and 7 complexes, respectively (Fig. 5A,5B,5C). GRASP also predicted a more diverse set of structures for these complexes (8/14 with more than one unique structure) than Boltz2 (4/7) and AlphaLink2 (0/6) (Supp Fig. 5A).

**Figure 5:**
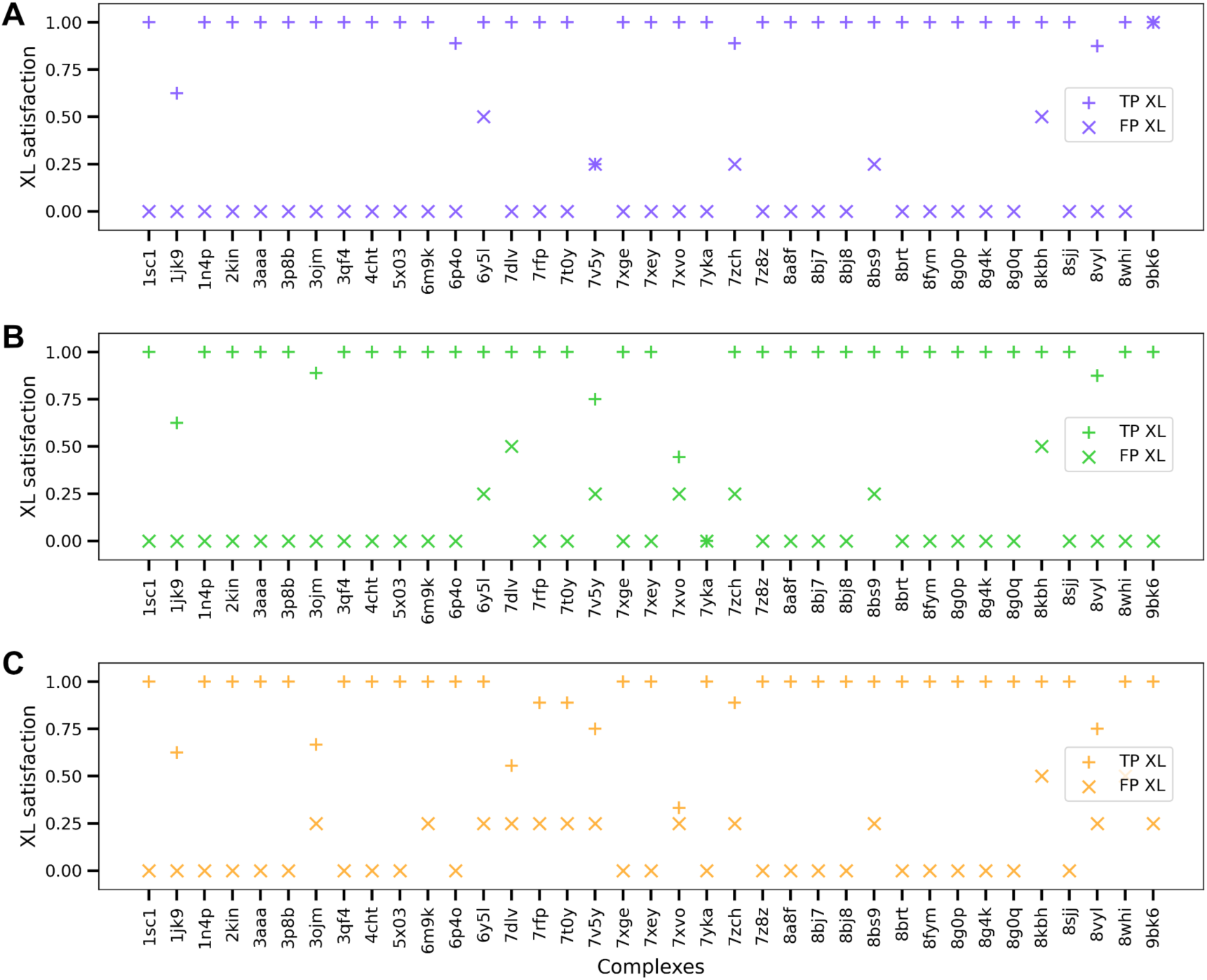
**Effect of false-positives**. For each complex, (A-C) the fraction of true positive (TP) and false positive (FP) crosslink (XL) satisfaction is shown for AlphaLink2 (purple), Boltz2 (green), and GRASP (orange).

Further, the predictions for most complexes do not change as the fraction of FP XLs increases. Specifically, at least one structure differed (DockQ < 0.23) between the two cases (10% FP, 33% FP) for 4/37, 9/37, and 16/37 complexes for AlphaLink2, Boltz2, and GRASP, respectively. In principle, satisfying a higher fraction of adversarial FP crosslinks would require sampling conformations that differ substantially from those that satisfy the TP XLs. However, the methods show little evidence of sampling alternative conformations. Among the methods, GRASP appears to be the most responsive to crosslinks guidance, generating alternative conformations for several complexes. Finally, the stereochemical quality of the predicted structures does not appear to be compromised by the addition of FP crosslinks, as measured by the mean clashscore and mean Molprobity score (Supp Fig. 5B,5C).

### Does data guidance allow predicting multiple states?

The crosslink datasets used so far were derived from a single native structure per complex. As a result, it is unclear whether the limited diversity of the predicted structures reflects a limitation of the AF-based IM methods or simply the lack of alternative structures consistent with the crosslinks. We next examined the ability of these methods to predict multiple states from crosslinks. In the absence of experimental structures for multi-state protein complexes, we evaluated two monomeric proteins known to exist in apo and holo conformations ^26^.

Ribonuclease H (RNase H) is an endonuclease enzyme that degrades RNA in RNA- DNA hybrids. Calmodulin1 is part of the calcium signalling pathway and regulates various enzymes and ion channels. The TM-score between the apo and holo states for RNaseH (Calmodulin1) is 0.33 (0.43). We sought to determine whether the AF- based methods predict both the apo and holo state structures, given a combined set of crosslinks derived from both states. Additionally, we investigated whether the predictions could be guided towards a specific state using crosslinks derived exclusively from either the apo or holo state (See Methods). We assessed the performance of each method using the TM-score of the predicted structures to the apo and holo state structures.

For both proteins, the predictions from all three methods were biased towards the holo state, regardless of the source of crosslinks (Fig. 6, 7, Suppl Fig. 6). AlphaLink2 and GRASP were unable to predict the apo state structure for both proteins, even when provided crosslinks derived exclusively from the apo state as input. Boltz2 predicted a few structures similar to the apo state for Calmodulin1. However, its predictions, like those of AlphaLink2 and GRASP, appear to be unaffected by the source of the input crosslinks (Fig. 6).

**Figure 6:**
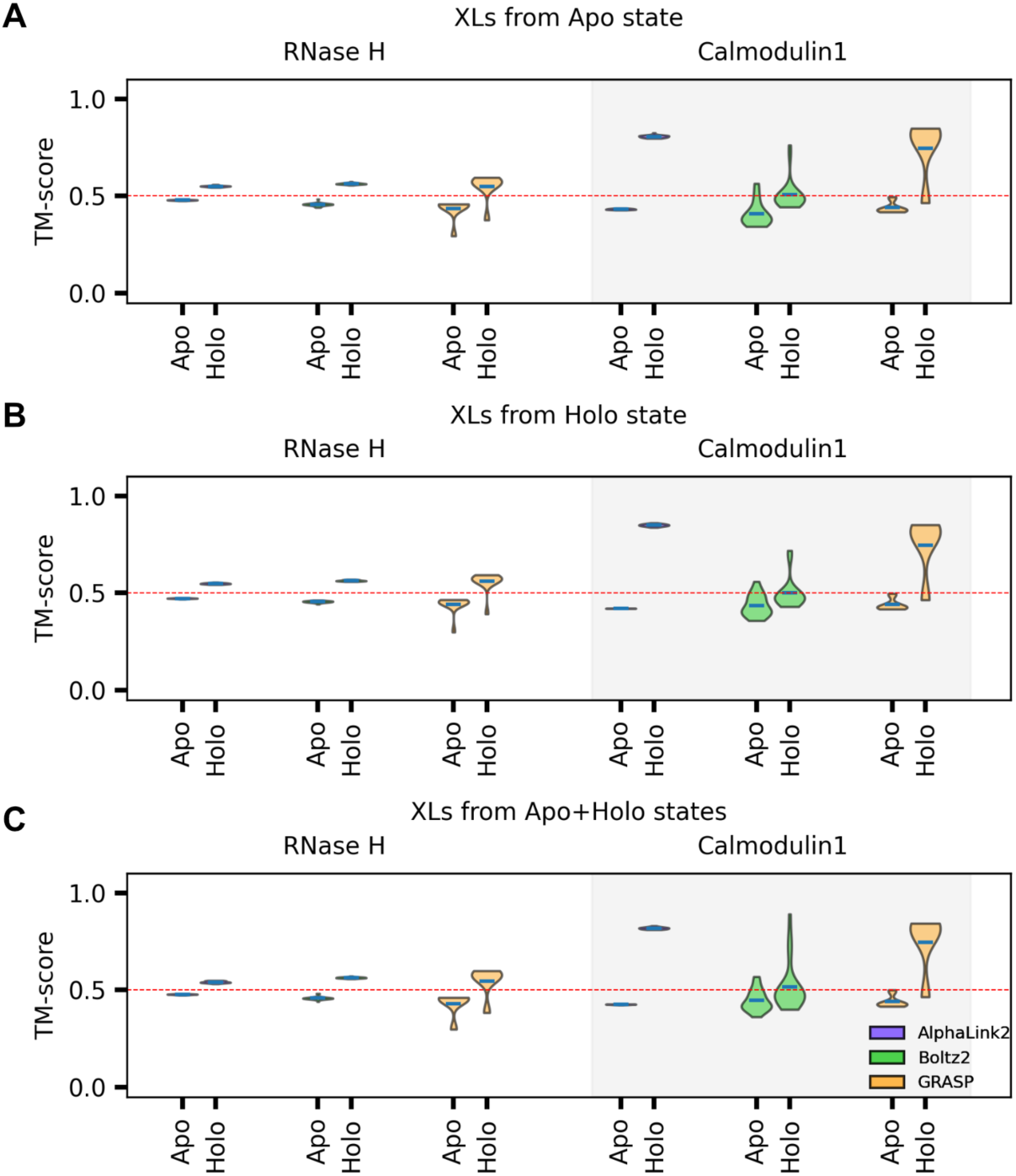
**Assessing exploration of multiple states**. The distribution of TM-score for the set of predicted structures with respect to the apo and holo states is shown for input crosslinks derived from (A) the apo state, (B) the holo state, and (C) the apo and holo states combined.

**Figure 7:**
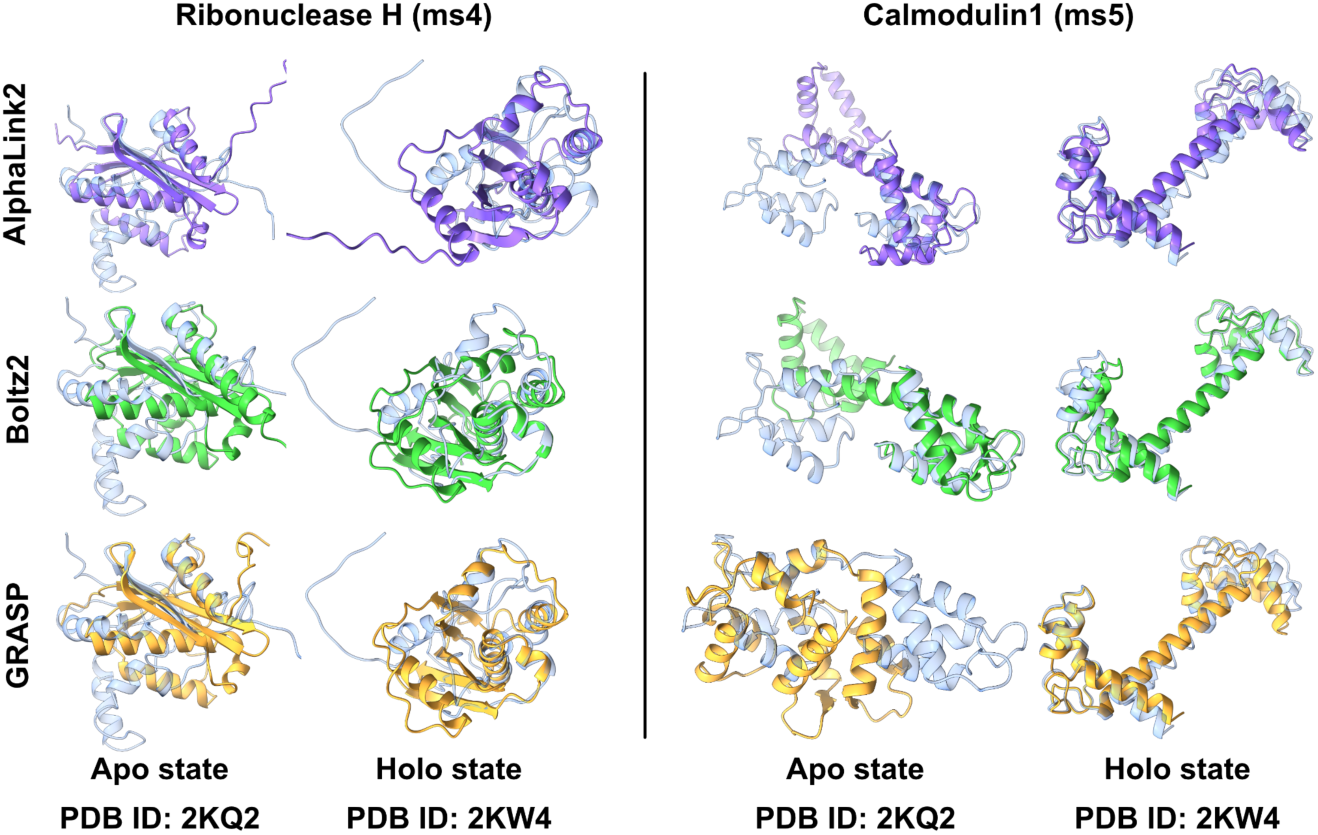
**Predicted structures for multi-state proteins**. For two examples of multi- state proteins, the predicted structures (solid colours) with the highest TM-scores relative to the apo and holo states (transparent) are shown for each method.

Besides, the distribution of the TM-score for the predicted structures is narrow, especially for AlphaLink2 (Fig. 6). Overall, the performance of these methods indicates insufficient sampling of alternative structures, plausibly due to training bias. Consequently, crosslinks guidance may be insufficient to overcome these learned biases and drive predictions towards configurations that lie outside the model’s learned structural prior.

## Discussion

The ability to incorporate experimental data into structure predictors like AlphaFold has opened new avenues for integrative modeling of macromolecular complexes. However, it is unclear how these methods balance the input data with AlphaFold’s learned structural priors. Here, we evaluated three methods, AlphaLink2, Boltz2, and GRASP, on a benchmark of homomeric and heteromeric protein complexes, using chemical crosslinks as input data. On a dataset of mostly true positive (TP) crosslinks obtained from a single experimental structure, all three methods predicted structures with high crosslink satisfaction. These structures were also accurate, as measured by the DockQ score to the native structure and the MolProbity score. Of the three methods, Boltz2 was the fastest, whereas AlphaLink2 required the most computational time (Supp Fig. 6).

Yet, these methods possess significant limitations for integrative modeling. Current methods are not scalable for modeling large macromolecular assemblies, as evidenced by the limitation to obtain predictions for complexes with more than 1400 residues on a high-memory GPU ^13^. Moreover, the examined methods are not sufficiently sensitive to crosslinks guidance. For most complexes, the methods produce nearly identical predictions with and without the input crosslinks (Suppl Fig. 2). Further, the predictions remain largely unchanged as the fraction of false-positive crosslinks increases, even though satisfying these crosslinks would require predicting more diverse structures. Finally, when challenged with input crosslinks from distinct apo and holo states, the methods remain biased toward one of the states (Fig. 6).

This insensitivity may arise because AlphaFold’s learned structural priors may dominate the input data, potentially biasing the predictions towards certain conformations. As a result, this limits exploration of alternate conformations consistent with the data. However, sampling diverse conformations is crucial for integrative modeling as multiple structures may fit the potentially sparse, noisy, ambiguous, and heterogeneous input data ^3^. Overall, our study exposes limitations of Alphafold-based integrative modeling approaches: their inability to appropriately balance experimental evidence with model priors, and their failure to account for the structural heterogeneity that characterizes many systems. Future methods that address these shortcomings will be essential for integrative modeling.

## Methods

### Dataset

#### Complex benchmark

We created a benchmark dataset comprising multimeric complexes non-redundant with the AlphaFold training set obtained from the PDB benchmark from Mirabello C. *et al.* (2024), FoldBench, PINDER-XL, AFM benchmark, and SAbDAb (Fig. 1)^27–31^. From FoldBench, we selected protein-protein, protein-peptide, and antigen-antibody complexes. From SAbDab, we selected antibody structures that were non-redundant at 60% sequence identity and had a resolution lower than 4 Å. This resulted in a total of 2804 structures of complexes. From these structures, chain termini were trimmed to remove missing residues. Further, we removed complexes larger than 1400 residues, those containing non-standard amino acids, and those containing chains with more than 1% missing residues.

#### Simulating crosslinks

Next, we simulated crosslinks using JWalk ^32^. We selected all inter-protein Lys-Lys crosslinks from JWalk with a solvent-accessible surface distance (SASD) less than 20 Å. Complexes with fewer than eight inter-protein crosslinks were removed. For complexes with more than eight inter-protein crosslinks, a random subset of eight crosslinks was selected as true positives (TP). Additionally, we added 10% adversarial false-positive (FP) crosslinks, *i.e.,* one FP crosslink (1 FP XL) per complex with a SASD of greater than 40 Å. With this, our 10% FP dataset contained nine crosslinks per complex (8 TP and 1 FP). We also created a second dataset containing a higher fraction of FP crosslinks to simulate greater noise. For each complex, we added three FP crosslinks to the existing nine crosslinks to create a dataset with 33% FP crosslinks. Longer crosslinks (>20 Å SASD) were not selected since several AlphaFold-based methods are only sensitive to distances smaller than 22 Å.

#### Multi-state proteins

Due to the lack of multi-state protein complexes with fewer than 1400 residues, we use monomeric proteins known to exist in apo and holo states. We selected two such proteins from an existing multi-state protein benchmark – Ribonuclease H (PDB IDs: 2KQ2 and 2KW4) and Calmodulin1 (PDB IDs: 1DMO and 3CLN) ^26^. Of the 91 proteins in the benchmark, our selection was based on the following two criteria: (a) the TM- score between the apo and holo states must be lower than 0.5, (b) at least five mutually exclusive, state-specific crosslinks must be obtained for each of the states. We considered all 20 standard amino acids for computing crosslinks from JWalk within a maximum SASD of 20 Å. For each protein, we considered the maximum contiguous sequence in the PDB entries for the apo and holo states.

#### Predictions from the AlphaFold-based Integrative Modeling methods

For this study, we selected three AF-based methods that allow incorporating residue- level interaction data as obtained from XLMS and enable predictions for multimeric complexes: (i) AlphaLink2, (ii) Boltz2, and (iii) GRASP ^18,20,21^. To account for the variability in the predictions arising from differences in the input MSAs, we used the OpenFold MSA creation pipeline to create MSAs for all complexes in the benchmark^33^. Complexes for which the MSA creation failed were excluded. The predictions were obtained using the default arguments for all three methods, unless stated otherwise in the paper. A total of 25 structures were predicted by each method. For three complexes (5XCT, 6IWW, 7AGF), we were unable to obtain predictions from AlphaLink2 and Boltz2, and hence these were excluded. Thus, a total of 37 complexes were considered for evaluation.

### Data satisfaction

We compute the fraction of crosslinks satisfied by each structure in the predicted set of structures. A crosslink is considered satisfied if the Cα-Cα distance between the crosslinked residues is smaller than the maximum allowed crosslink distance of 20 Å. We account for ambiguity by considering a crosslink satisfied if any of the multiple copies of a protein in the complex satisfy the crosslink. Thus, the crosslink satisfaction for a given predicted structure, *S_xl_* is computed as,

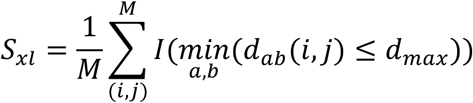

where, *M* represents the set of all crosslink residue pairs (*i*, *j*), *d_max_* represents the maximum allowed crosslink distance, *d_ab_*(*i*, *j*) represents the distance between residue *i* in copy *a* and residue *j* in copy *b*, and *I* is the indicator function. For each complex, we considered the maximum crosslink satisfaction across the predicted set of structures.

### Structural accuracy

For assessing structural accuracy, we computed the TM-score and DockQ score ^24,34^. To assess the structural accuracy of complexes, we measured the interface similarity of the predicted structures to the native structure, using the DockQ score, computed *via* the DockQ-v2 Python library ^24^. We evaluated methods based on the maximum DockQ score across their set of predicted structures for each complex. For multi-state proteins, we used the TM-score, as computed from USalign, to estimate the accuracy of each predicted structure with respect to the native apo and holo state structures ^34^.

### Stereochemical plausibility

We used MolProbity from the Phenix suite for measuring stereochemical plausibility^25^. We computed the MolProbity score and interatomic clashscore for all predicted structures. We evaluated the set of predicted structures from each method by their mean MolProbity score and mean clashscore.

### Structural diversity

We measured diversity in the set of 25 predicted structures per complex per method by determining the number of structurally distinct interfaces. We computed all-vs-all pairwise DockQ scores to evaluate interface similarity between all pairs of structures, considering two structures as similar if their DockQ score exceeded 0.23 ^23,24^. We then iteratively eliminated structurally similar predictions (DockQ > 0.23) to obtain a non- redundant set of structures.

## Data Availability

The datasets created in this manuscript have been deposited to Zenodo at https://doi.org/10.5281/zenodo.22159054.

The deposition contains the benchmark dataset used in the study, the predictions from Boltz2 and GRASP, and the figures presented in the paper.

The scripts to create the benchmark dataset, obtain predictions from AlphaLink2, Boltz2, and GRASP, and perform analysis are available at GitHub https://github.com/isblab/af_im.

Any additional information required to reanalyze the data reported in this paper is available from the lead contact upon request.

## Acknowledgement

We thank ISB Lab members Muskaan Jindal, Omkar Golatkar, and Mubashira KP for their useful comments on the manuscript. Molecular graphics images were produced using the UCSF ChimeraX packages from the Resource for Biocomputing, Visualization, and Informatics at the University of California, San Francisco (supported by NIH P41 RR001081, NIH R01-GM129325, and National Institute of Allergy and Infectious Diseases).

## Author contribution

Conceptualization: K.M., S.V. Reading and synthesis: K.M., S.V. Data curation: K.M.

Methodology: K.M., S.V. Writing – original draft: K.M.

Writing – review & editing: K.M., S.V.

Resources, Supervision, Funding acquisition: S.V.

## Funding

This work has been supported by the following grants: Department of Atomic Energy (DAE) TIFR grant RTI 4006 and 4018, and Department of Biotechnology (DBT) grant BT/PR40323/BTIS/137/78/2023 from the Government of India to S.V. and the “HS Gupta and Asha Gupta Research Travel Fellowship" by Centre for Artificial Learning & Intelligence for Biological Research & Education (CALIBRE) at NCBS to K.M.

## Declaration of interests

None declared.

## Declaration of generative AI and AI-assisted technologies

During the preparation of this work, the authors used ChatGPT 5.6 (Aug-Sep 2026) to refine the text from initial drafts. After using these tools, the authors reviewed and edited the content as needed and take full responsibility for the content of the publication.

## Notes

https://doi.org/10.5281/zenodo.22159054

### Competing Interest Statement

The authors have declared no competing interest.

### Summary of Updates

The abstract was revised. The subsection title "Robustness to noise" was changed to "Effect of false-positive crosslinks" in the Results section, and its contents were revised. Discussion section was revised.

