## Supplementary material for "Evaluation of methods for AlphaFold-based integrative modeling"

### Supplementary figures

**Supp Figure 1: Maximum crosslink satisfaction for guided and unguided predictions**. For Boltz2 (green, triangle) and GRASP (orange, circle), we show the maximum crosslink satisfaction achieved by a structure in the set of predicted structures for each complex, comparing guided vs unguided predictions.

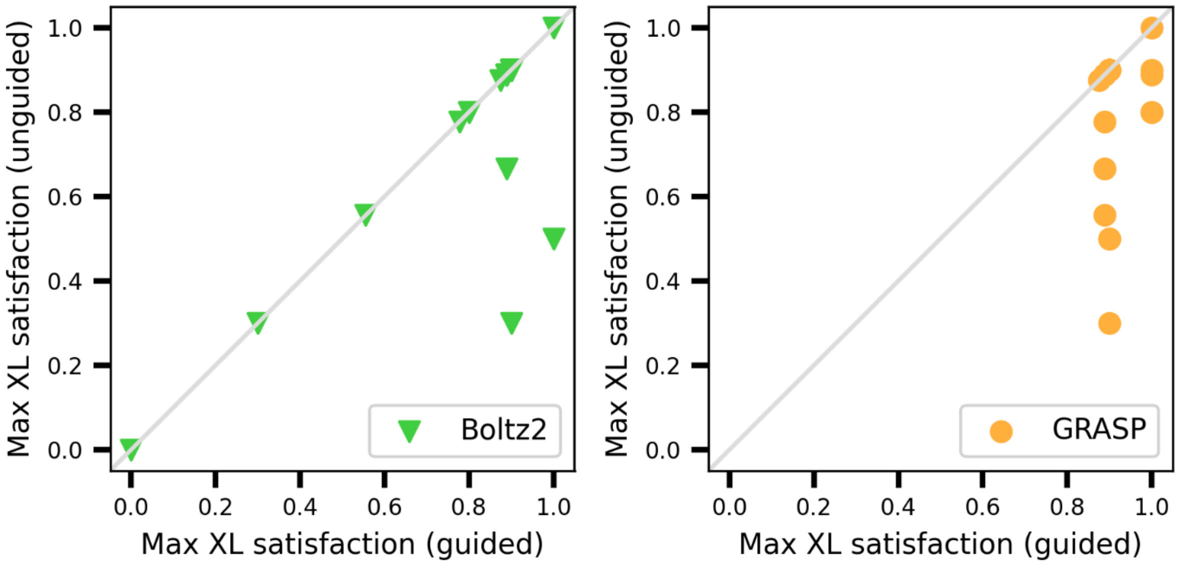

**Supp Figure 2: Distribution of crosslink satisfaction for guided and unguided predictions**. The distribution of crosslink satisfaction in the predicted set for each complex, comparing restraint-guided vs unguided predictions for (A) Boltz2 and (B) GRASP.

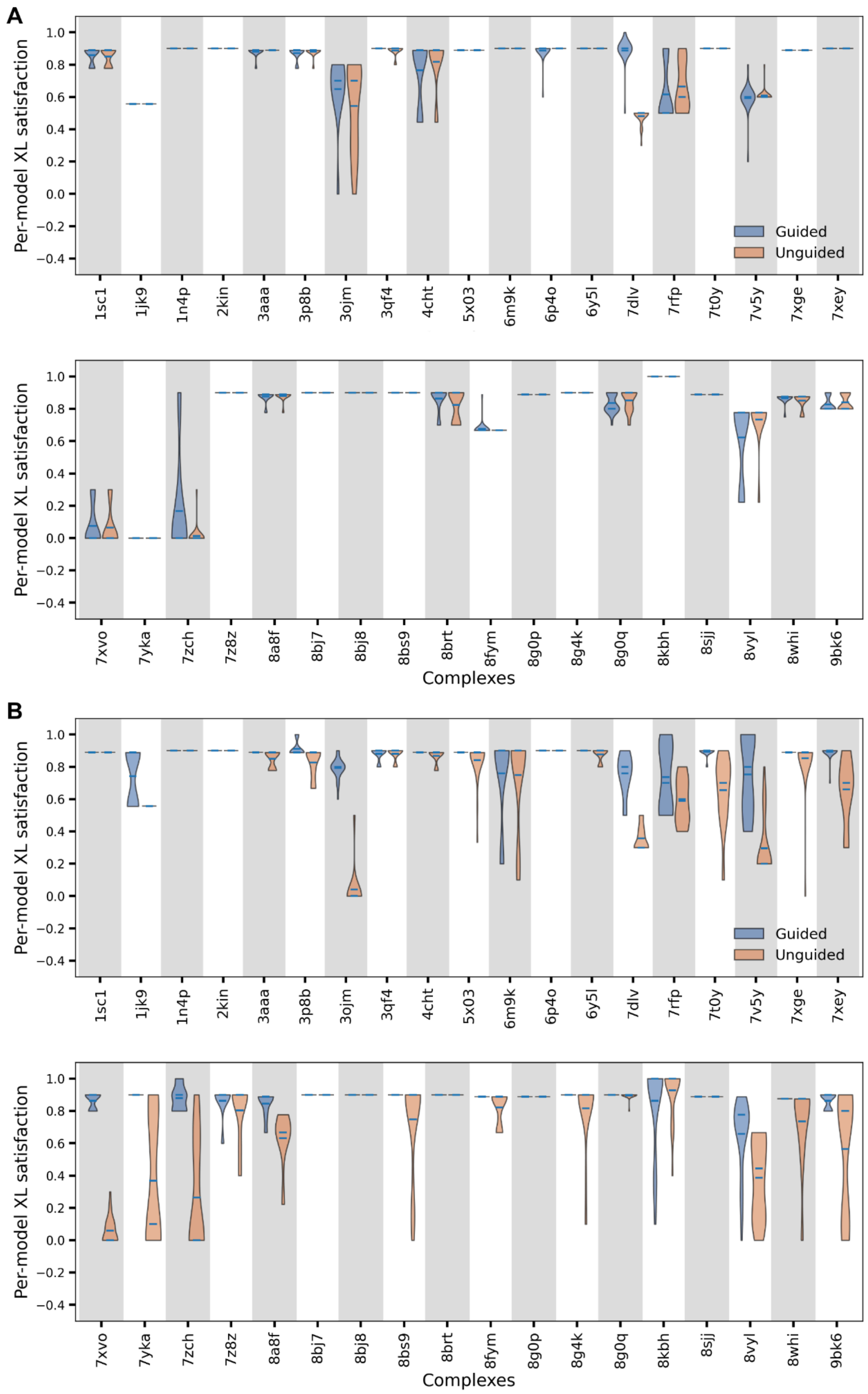

**Supp Figure 3: Assessing mean clashscore**. The mean clashscore of the predicted set of structures vs the clashscore for the native structure for each complex for AlphaLink2 (purple, square), Boltz2 (green, triangle), and GRASP (orange, circle) is shown.

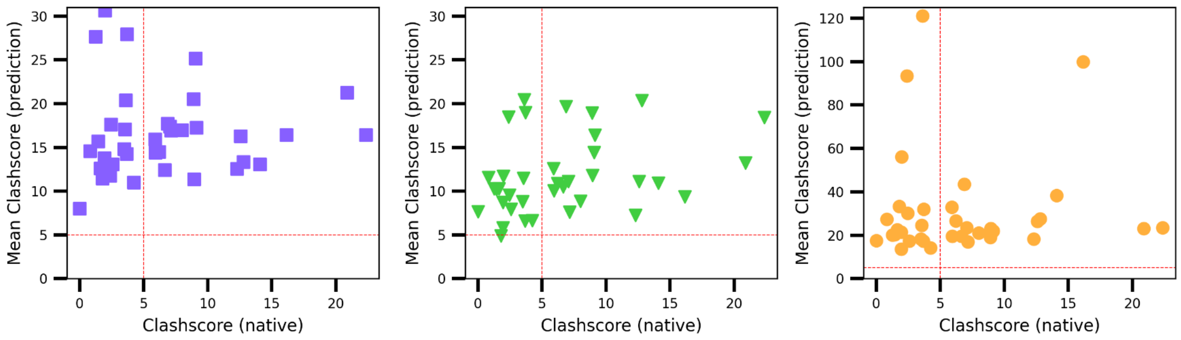

**Supp Figure 4: Assessing predicted model confidence**. The mean model confidence across the predicted set of structures by AlphaLink2 (purple, square), Boltz2 (green, triangle), and GRASP (orange, circle) for each complex is shown.

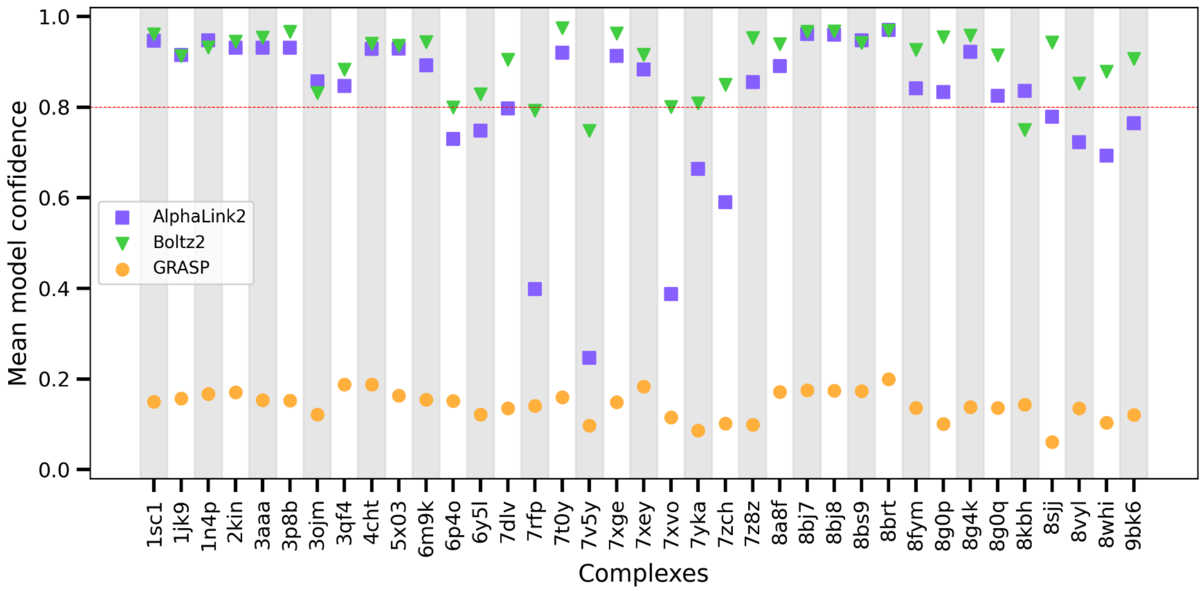

**Supp Figure 5: Effect of false positive crosslinks**. For the predictions obtained from AlphaLink2 (purple, square), Boltz2 (green, triangle), and GRASP (orange, circle) using 10% vs 33% false positive (FP) crosslinks, (A) the number of unique predicted structures based on DockQ and (B) the mean molProbity score are shown.

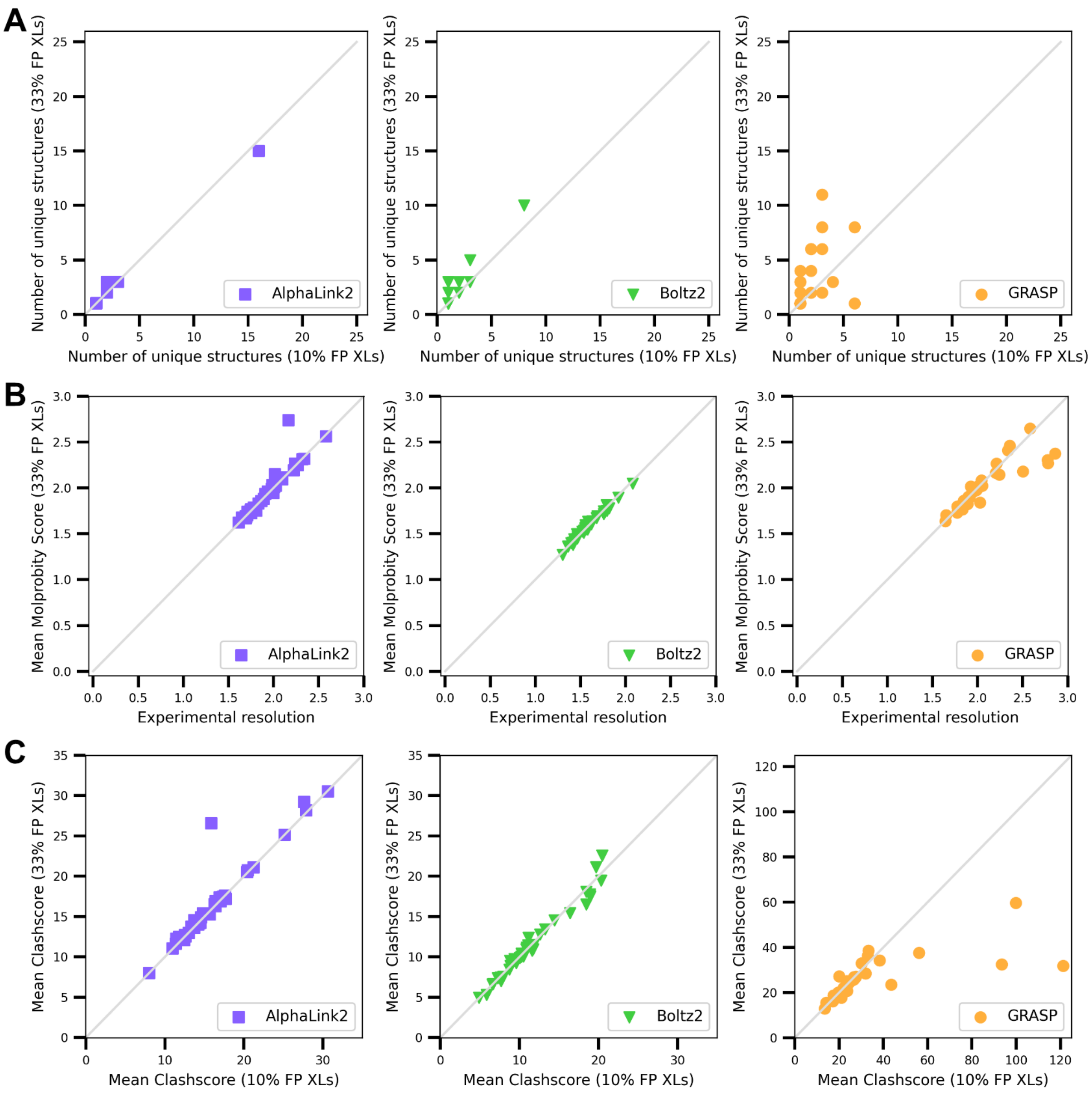

**Supp Figure 7: Time taken for predictions**. We measure the mean time taken (in hours) by each method across the benchmark. AlphaLink2 (purple), Boltz2 (green), and GRASP (orange). Runtimes were computed on an AMD EPYC 7763 64-Core Processor, with 1024 GB RAM, and an A6000 GPU.

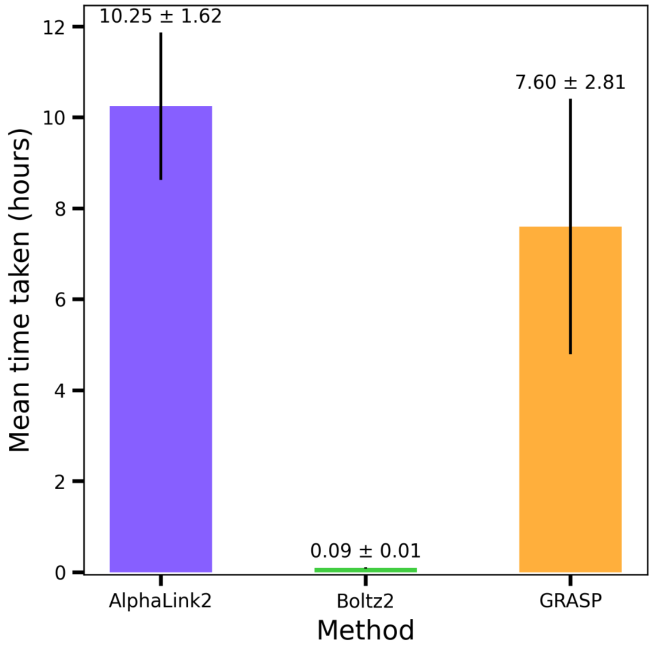

### Supplementary tables

**Supp Table 1: Assessing the performance of AlphaFold-based integrative modeling methods**. For each complex in the benchmark, the crosslink satisfaction, DockQ, number of unique structures, Molprobity score, and clashscore are reported for AlphaLink2, Boltz2, and GRASP.

| Complex | Method | Crosslink satisfaction | | DockQ | | Number of structures with unique interface | Mean Molprobity score | Mean clashscore |
|---|---|---|---|---|---|---|---|---|
| | | Max | Mean | Max | Mean | | | |
| **1sc1** | Alphalink2 | 0.889 | 0.889 | 0.6 | 0.6 | 1 | 1.864 | 14.396 |
| | Boltz2 | 0.889 | 0.858 | 0.5 | 0.5 | 1 | 1.618 | 10.032 |
| | Grasp | 0.889 | 0.889 | 0.6 | 0.6 | 1 | 1.99 | 19.636 |
| **1jk9** | Alphalink2 | 0.556 | 0.556 | 0.5 | 0.404 | 1 | 1.78 | 16.398 |
| | Boltz2 | 0.556 | 0.556 | 0.5 | 0.452 | 1 | 1.758 | 18.446 |
| | Grasp | 0.889 | 0.742 | 0.5 | 0.44 | 1 | 2.028 | 23.556 |
| **1n4p** | Alphalink2 | 0.9 | 0.9 | 0.9 | 0.9 | 1 | 1.893 | 17.226 |
| | Boltz2 | 0.9 | 0.9 | 0.8 | 0.62 | 1 | 1.919 | 16.393 |
| | Grasp | 0.9 | 0.9 | 0.9 | 0.836 | 1 | 1.956 | 22.031 |
| **2kin** | Alphalink2 | 0.9 | 0.9 | 0.8 | 0.748 | 1 | 1.694 | 12.532 |
| | Boltz2 | 0.9 | 0.9 | 0.8 | 0.728 | 1 | 1.397 | 7.267 |
| | Grasp | 0.9 | 0.9 | 0.8 | 0.708 | 1 | 1.883 | 18.3 |
| **3aaa** | Alphalink2 | 0.889 | 0.787 | 0.7 | 0.7 | 1 | 1.711 | 14.23 |
| | Boltz2 | 0.889 | 0.88 | 0.7 | 0.7 | 1 | 1.361 | 6.585 |

| | | | | | | | | |
|---|---|---|---|---|---|---|---|---|
| | Grasp | 0.889 | 0.889 | 0.8 | 0.8 | 1 | 1.772 | 17.386 |
| **3p8b** | Alphalink2 | 0.889 | 0.889 | 0.8 | 0.708 | 1 | 2.008 | 16.965 |
| | Boltz2 | 0.889 | 0.871 | 0.9 | 0.752 | 1 | 1.47 | 8.839 |
| | Grasp | 1 | 0.911 | 0.9 | 0.664 | 1 | 1.833 | 21.093 |
| **3ojm** | Alphalink2 | 0.9 | 0.804 | 0.7 | 0.544 | 1 | 2.072 | 13.335 |
| | Boltz2 | 0.8 | 0.648 | 0.3 | 0.152 | 3 | 1.817 | 20.35 |
| | Grasp | 0.9 | 0.792 | 0.7 | 0.268 | 6 | 2.211 | 27.55 |
| **3qf4** | Alphalink2 | 0.9 | 0.9 | 0.8 | 0.8 | 1 | 2.583 | 21.22 |
| | Boltz2 | 0.9 | 0.9 | 0.4 | 0.4 | 1 | 1.629 | 13.238 |
| | Grasp | 0.9 | 0.88 | 0.7 | 0.56 | 1 | 2.012 | 23.052 |
| **4cht** | Alphalink2 | 0.889 | 0.889 | 0.8 | 0.796 | 1 | 1.909 | 13.041 |
| | Boltz2 | 0.889 | 0.764 | 0.8 | 0.744 | 1 | 1.432 | 7.946 |
| | Grasp | 0.889 | 0.889 | 0.8 | 0.712 | 1 | 1.845 | 17.428 |
| **5x03** | Alphalink2 | 0.889 | 0.889 | 0.7 | 0.7 | 1 | 1.751 | 11.365 |
| | Boltz2 | 0.889 | 0.889 | 0.7 | 0.688 | 1 | 1.586 | 11.783 |
| | Grasp | 0.889 | 0.889 | 0.7 | 0.684 | 1 | 1.847 | 23.058 |
| **6m9k** | Alphalink2 | 0.9 | 0.9 | 0.8 | 0.76 | 1 | 1.697 | 15.667 |
| | Boltz2 | 0.9 | 0.9 | 0.8 | 0.8 | 1 | 1.532 | 10.206 |
| | Grasp | 0.9 | 0.76 | 0.8 | 0.616 | 2 | 1.796 | 20.232 |
| **6p4o** | Alphalink2 | 0.8 | 0.68 | 0.2 | 0.2 | 1 | 2.319 | 20.489 |
| | Boltz2 | 0.9 | 0.888 | 0.2 | 0.136 | 1 | 1.793 | 18.926 |
| | Grasp | 0.9 | 0.9 | 0.3 | 0.3 | 1 | 1.986 | 19.01 |
| **6y5l** | Alphalink2 | 0.9 | 0.828 | 0.4 | 0.272 | 1 | 2.165 | 15.884 |

| | | | | | | | | |
|---|---|---|---|---|---|---|---|---|
| | Boltz2 | 0.9 | 0.9 | 0.4 | 0.296 | 1 | 1.583 | 12.601 |
| | Grasp | 0.9 | 0.9 | 0.1 | 0.076 | 1 | 2.338 | 32.886 |
| **7dlv** | Alphalink2 | 0.9 | 0.868 | 0.8 | 0.684 | 1 | 1.782 | 17.695 |
| | Boltz2 | 1 | 0.888 | 0.7 | 0.648 | 2 | 1.789 | 19.67 |
| | Grasp | 0.9 | 0.76 | 0.6 | 0.328 | 6 | 2.503 | 43.497 |
| **7rfp** | Alphalink2 | 0.8 | 0.58 | 0.3 | 0.128 | 2 | 2.267 | 20.384 |
| | Boltz2 | 0.9 | 0.616 | 0.4 | 0.284 | 1 | 2.08 | 20.478 |
| | Grasp | 1 | 0.736 | 0.3 | 0.132 | 3 | 2.781 | 121.186 |
| **7t0y** | Alphalink2 | 0.9 | 0.9 | 0.9 | 0.78 | 1 | 2.095 | 11.449 |
| | Boltz2 | 0.9 | 0.9 | 0.9 | 0.872 | 1 | 1.429 | 4.883 |
| | Grasp | 0.9 | 0.892 | 0.3 | 0.188 | 1 | 2.58 | 33.296 |
| **7v5y** | Alphalink2 | 0.2 | 0.2 | 0.8 | 0.72 | 1 | 1.755 | 11.778 |
| | Boltz2 | 0.8 | 0.592 | 0.3 | 0.196 | 1 | 1.756 | 18.458 |
| | Grasp | 1 | 0.752 | 0.3 | 0.124 | 3 | 2.86 | 93.426 |
| **7xge** | Alphalink2 | 0.889 | 0.889 | 0.9 | 0.84 | 1 | 2.238 | 16.9 |
| | Boltz2 | 0.889 | 0.889 | 0.9 | 0.9 | 1 | 1.413 | 7.634 |
| | Grasp | 0.889 | 0.889 | 0.9 | 0.88 | 1 | 1.774 | 17.034 |
| **7xey** | Alphalink2 | 0.9 | 0.848 | 0.7 | 0.588 | 1 | 2.224 | 17.394 |
| | Boltz2 | 0.9 | 0.9 | 0.8 | 0.8 | 1 | 1.56 | 11.108 |
| | Grasp | 0.9 | 0.892 | 0.6 | 0.56 | 1 | 2.052 | 23.438 |
| **7xvo** | Alphalink2 | 0.9 | 0.36 | 0.7 | 0.184 | 16 | 1.81 | 12.414 |
| | Boltz2 | 0.3 | 0.076 | 0 | 0 | 8 | 1.536 | 10.496 |
| | Grasp | 0.9 | 0.864 | 0.7 | 0.408 | 3 | 1.94 | 19.55 |

| | | | | | | | | |
|---|---|---|---|---|---|---|---|---|
| **7yka** | Alphalink2 | 0.9 | 0.9 | 0.3 | 0.3 | 1 | 1.936 | 27.625 |
| | Boltz2 | 0 | 0 | 0 | 0 | 3 | 1.521 | 10.266 |
| | Grasp | 0.9 | 0.9 | 0.8 | 0.264 | 2 | 1.924 | 20.075 |
| **7zch** | Alphalink2 | 0.8 | 0.728 | 0.4 | 0.26 | 3 | 2.017 | 16.43 |
| | Boltz2 | 0.9 | 0.168 | 0.5 | 0.072 | 2 | 1.473 | 9.356 |
| | Grasp | 1 | 0.88 | 0.4 | 0.184 | 2 | 2.776 | 99.886 |
| **7z8z** | Alphalink2 | 0.9 | 0.9 | 0.8 | 0.72 | 1 | 2.027 | 27.904 |
| | Boltz2 | 0.9 | 0.9 | 0.8 | 0.716 | 1 | 1.771 | 18.976 |
| | Grasp | 0.9 | 0.864 | 0.8 | 0.704 | 1 | 2.001 | 32.012 |
| **8a8f** | Alphalink2 | 0.889 | 0.889 | 0.6 | 0.6 | 1 | 2.1 | 17.624 |
| | Boltz2 | 0.889 | 0.876 | 0.6 | 0.556 | 1 | 1.679 | 9.568 |
| | Grasp | 0.889 | 0.844 | 0.6 | 0.444 | 2 | 2.356 | 30.17 |
| **8bj7** | Alphalink2 | 0.9 | 0.9 | 0.9 | 0.9 | 1 | 1.762 | 14.481 |
| | Boltz2 | 0.9 | 0.9 | 0.9 | 0.9 | 1 | 1.553 | 10.865 |
| | Grasp | 0.9 | 0.9 | 0.9 | 0.796 | 1 | 1.898 | 26.693 |
| **8bj8** | Alphalink2 | 0.9 | 0.9 | 0.9 | 0.9 | 1 | 1.755 | 14.58 |
| | Boltz2 | 0.9 | 0.9 | 0.9 | 0.9 | 1 | 1.578 | 11.591 |
| | Grasp | 0.9 | 0.9 | 0.9 | 0.8 | 1 | 1.914 | 27.328 |
| **8bs9** | Alphalink2 | 0.9 | 0.9 | 0.9 | 0.9 | 1 | 1.689 | 13.715 |
| | Boltz2 | 0.9 | 0.9 | 0.9 | 0.812 | 1 | 1.465 | 8.71 |
| | Grasp | 0.9 | 0.9 | 0.8 | 0.716 | 1 | 1.888 | 21.472 |
| **8brt** | Alphalink2 | 0.9 | 0.9 | 0.9 | 0.9 | 1 | 1.744 | 12.608 |
| | Boltz2 | 0.9 | 0.864 | 0.9 | 0.9 | 1 | 1.541 | 10.288 |

| | | | | | | | | |
|---|---|---|---|---|---|---|---|---|
| | Grasp | 0.9 | 0.9 | 0.9 | 0.88 | 1 | 1.989 | 22.54 |
| **8fym** | Alphalink2 | 0.889 | 0.889 | 0.4 | 0.324 | 1 | 1.834 | 17.05 |
| | Boltz2 | 0.889 | 0.676 | 0.3 | 0.232 | 1 | 1.572 | 11.497 |
| | Grasp | 0.889 | 0.889 | 0.6 | 0.392 | 1 | 2.004 | 24.519 |
| **8g0p** | Alphalink2 | 0.889 | 0.889 | 0.6 | 0.6 | 1 | 1.617 | 10.982 |
| | Boltz2 | 0.889 | 0.889 | 0.9 | 0.88 | 1 | 1.363 | 6.668 |
| | Grasp | 0.889 | 0.889 | 0.8 | 0.684 | 1 | 1.65 | 14.156 |
| **8g4k** | Alphalink2 | 0.9 | 0.9 | 0.8 | 0.724 | 1 | 1.667 | 7.989 |
| | Boltz2 | 0.9 | 0.9 | 0.8 | 0.7 | 1 | 1.416 | 7.644 |
| | Grasp | 0.9 | 0.9 | 0.8 | 0.692 | 1 | 1.808 | 17.478 |
| **8g0q** | Alphalink2 | 0.9 | 0.9 | 0.6 | 0.58 | 1 | 2.342 | 25.167 |
| | Boltz2 | 0.9 | 0.836 | 0.6 | 0.556 | 1 | 1.685 | 14.437 |
| | Grasp | 0.9 | 0.9 | 0.6 | 0.544 | 1 | 2.201 | 21.624 |
| **8kbh** | Alphalink2 | 1 | 1 | 0.8 | 0.8 | 1 | 1.712 | 16.283 |
| | Boltz2 | 1 | 1 | 0.8 | 0.752 | 1 | 1.556 | 11.121 |
| | Grasp | 1 | 0.864 | 0.8 | 0.668 | 3 | 1.946 | 26.52 |
| **8sjj** | Alphalink2 | 0.889 | 0.889 | 0.7 | 0.7 | 1 | 1.65 | 13.733 |
| | Boltz2 | 0.889 | 0.889 | 0.8 | 0.7 | 1 | 1.302 | 5.83 |
| | Grasp | 0.889 | 0.889 | 0.8 | 0.688 | 1 | 1.641 | 13.586 |
| **8vyl** | Alphalink2 | 0.778 | 0.751 | 0.7 | 0.528 | 1 | 1.978 | 13.034 |
| | Boltz2 | 0.778 | 0.622 | 0.5 | 0.364 | 1 | 1.552 | 10.916 |
| | Grasp | 0.889 | 0.658 | 0.5 | 0.348 | 1 | 2.044 | 38.262 |
| **8whi** | Alphalink2 | 0.875 | 0.875 | 0.5 | 0.496 | 1 | 1.999 | 14.8 |

| | | | | | | | | |
|---|---|---|---|---|---|---|---|---|
| | Boltz2 | 0.875 | 0.865 | 0.5 | 0.328 | 1 | 1.461 | 8.794 |
| | Grasp | 0.875 | 0.875 | 0.5 | 0.436 | 1 | 1.852 | 18.33 |
| **9bk6** | Alphalink2 | 0.9 | 0.868 | 0.8 | 0.768 | 2 | 1.986 | 30.649 |
| | Boltz2 | 0.9 | 0.828 | 0.7 | 0.688 | 1 | 1.572 | 11.71 |
| | Grasp | 0.9 | 0.864 | 0.7 | 0.576 | 4 | 2.243 | 56.174 |
